# Characterizing the genetic basis of trait evolution in the Mexican cavefish

**DOI:** 10.1101/2021.12.16.472318

**Authors:** Camila Oliva, Nicole K. Hinz, Wayne Robinson, Alexys M. Barrett Thompson, Julianna Booth, Lina M. Crisostomo, Samantha Zanineli, Maureen Tanner, Evan Lloyd, Morgan O’Gorman, Brittnee McDole, Alexandra Paz, Rob Kozol, Elizabeth B. Brown, Johanna E. Kowalko, Yaouen Fily, Erik R. Duboue, Alex C. Keene

## Abstract

Evolution in response to a change in ecology often coincides with various morphological, physiological, and behavioral traits. For most organisms little is known about the genetic and functional relationship between evolutionarily derived traits, representing a critical gap in our understanding of adaptation The Mexican tetra, *Astyanax mexicanus*, consists of largely independent populations of fish that inhabit at least 30 caves in Northeast Mexico, and a surface fish population, that inhabits the rivers of Mexico and Southern Texas. The recent application of molecular genetic approaches combined with behavioral phenotyping have established A. mexicanus as a model for studying the evolution of complex traits. Cave populations of A. mexicanus are interfertile with surface populations and have evolved numerous traits including eye degeneration, insomnia, albinism and enhanced mechanosensory function. The interfertility of different populations from the same species provides a unique opportunity to define the genetic relationship between evolved traits and assess the co-evolution of behavioral and morphological traits with one another. To define the relationships between morphological and behavioral traits, we developed a pipeline to test individual fish for multiple traits. This pipeline confirmed differences in locomotor activity, prey capture, and startle reflex between surface and cavefish populations. To measure the relationship between traits, individual F2 hybrid fish were characterized for locomotor behavior, prey-capture behavior, startle reflex and morphological attributes. Analysis revealed an association between body length and slower escape reflex, suggesting a trade-off between increased size and predator avoidance in cavefish. Overall, there were few associations between individual behavioral traits, or behavioral and morphological traits, suggesting independent genetic changes underlie the evolution of behavioral and morphological traits. Taken together, this approach provides a novel system to identify genes that underlie naturally occurring genetic variation in morphological and behavioral traits.

## Introduction

Environmental changes often drive the evolution of morphological, behavioral, and physiological traits (Rose 2007). Often trait evolution involves complex changes in genetic architecture that include pleiotropic gene function, or traits that independently evolve in parallel (Stern 2013), yet systematically testing the evolutionary relationship between traits has been challenging. Subterranean environments present a unique opportunity to investigate the relationship between environment and trait evolution because the changes in environment, such as loss of light, are often well defined and present across independently evolved populations (Culver and Pipan 2009). In addition, in many cases, a closely related species or population remains in the surface environment, allowing for a direct comparison between species that inhabit different environments (Elliott 2016; Borowsky 2018). Finally, many traits associated with subterranean evolution including albinism, reduced eye size, lower metabolic rate, and loss of circadian rhythms, have evolved in distantly-related species in cavefish and subterranean mammals(Poulson 2001; Jeffery 2009; Tian *et al*. 2017). Therefore, investigating trait evolution in subterranean species has potential to uncover whether seemingly distinct traits are genetically linked and may have co-evolved.

The Mexican tetra *Astyanax mexicanus* is a leading model to study the evolution of complex traits (Keene *et al*. 2015; Jeffery 2020). The convergent evolution of cavefish from surface-like ancestors in geographically distinct cave environments produced two morphologically distinct forms of *A. mexicanus*. The first is a surface-dwelling form with fully developed eyes found in above-ground rivers and streams of northeast Mexico and parts of southern Texas, and the second includes at least 30 populations cave-dwelling forms, mostly found within the Sierra del Abra region of northeast Mexico (Mitchell *et al*. 1977; Jeffery 2001; Gross 2012). Genomic and geological data suggest cavefish populations are largely independent (with some admixture) providing the opportunity to test the repeatability of evolution (Mitchell *et al*. 1977; Strecker *et al*. 2004; Herman *et al*. 2018). Cave-dwelling forms have converged on distinct morphological traits, including albinism and eye loss (Jeffery 2020). In addition, cavefish evolved numerous behavioral changes including different prey capture, startle response, and increased locomotor activity (Duboué *et al*. 2011; Lloyd *et al*. 2018; Paz *et al*. 2020). Overall these changes are thought to be critical for foraging in the absence of visual cues (Yoshizawa 2015; Keene and Duboue 2018; McGaugh *et al*. 2020). Therefore, the robust phenotypic differences between surface fish and cavefish provide an opportunity to examine the relationship between the evolution of behaviors and morphological traits.

Cavefish and surface fish are interfertile, allowing for the generation of hybrid fish that can be used to assess whether shared or independent genetic architecture regulates seemingly distinct cave-like traits (Protas *et al*. 2006; Yoshizawa *et al*. 2012; O’Quin and McGaugh 2016). Quantitative trait loci (QTL) analyses for multiple traits, including lens and eye size, as well as a relationship between vibration attraction behavior and superficial neuromast number, support the notion that genetic pleiotropy may contribute to the evolution of multiple traits (Yoshizawa *et al*. 2012; Kowalko *et al*. 2013; McGaugh *et al*. 2014). In addition, numerous functional interactions have been identified, including interactions between eye loss and the expansion of the jaw and hypothalamus (Yamamoto *et al*. 2009; Pottin *et al*. 2011; Atukorala and Franz-Odendaal 2018). Studies have also found genetic interactions between albinism, elevated catecholamines, and increased locomotor activity (Bilandzija *et al*. 2013; Bilandžija *et al*. 2018). A later study found that mutation of the *oca2* gene causes sleep loss and increased locomotor activity (O’Gorman *et al*. 2021). Therefore, investigating many different cave-evolved traits in individual hybrids has potential to identify the degree to which evolved traits to one another.

Here, we generated a pipeline for analyzing behavior and morphology in individual fish. We applied this to analyze F2 hybrids between surface and Pachón cavefish, a highly troglomorphic population, and measured the relationship between traits. We systematically investigated the relationships between individual behaviors as well as the relationships between these behaviors and morphological traits. These studies suggest that behavioral and morphological traits are largely regulated independently, suggesting independent evolution of many cave-associated traits.

## Results

In zebrafish and *A. mexicanus*, 6 days post-fertilization (dpf) larvae are often studied because their transparency and small size is amenable to brain imaging and high-throughput behavioral analysis (Halpern *et al*. 2008; Keene and Appelbaum 2019). We designed our experiments to measure behavior in 6 dpf fish, followed by morphological analysis at 7 dpf. To define morphological differences between surface and cavefish populations, we compared multiple anatomical traits in surface and Pachón cavefish. At this timepoint, the overall developmental stage of surface and cavefish are largely similar, allowing for direct comparisons of anatomical features (Hinaux *et al*. 2011). We quantified eight traits related to overall body size, craniofacial development, and pigmentation (Figure 1, A, B and Figure S1). At 7 dpf, cavefish were on average larger with increased body length, height, and head length, revealing an overall increase in the body and head size of Pachón cavefish (Figure S2 and Figure 1E). Consistent with the previous reports, we found that jaw size is significantly increased in cavefish and eye size is significantly reduced, raising the possibility of a trade-off between eye and jaw size (Figure 1C,D; Yamamoto et al. 2009; Pottin et al. 2011). Nearly all differences were maintained when individual traits were normalized to length body length (Figure S3), confirming that the observed differences are not due to overall changes in size. Together these analyses confirm the presence of numerous morphological differences between surface and Pachón cavefish at 7 dpf, providing a platform to investigate the genetic relationship between these traits.

**Figure 1.**
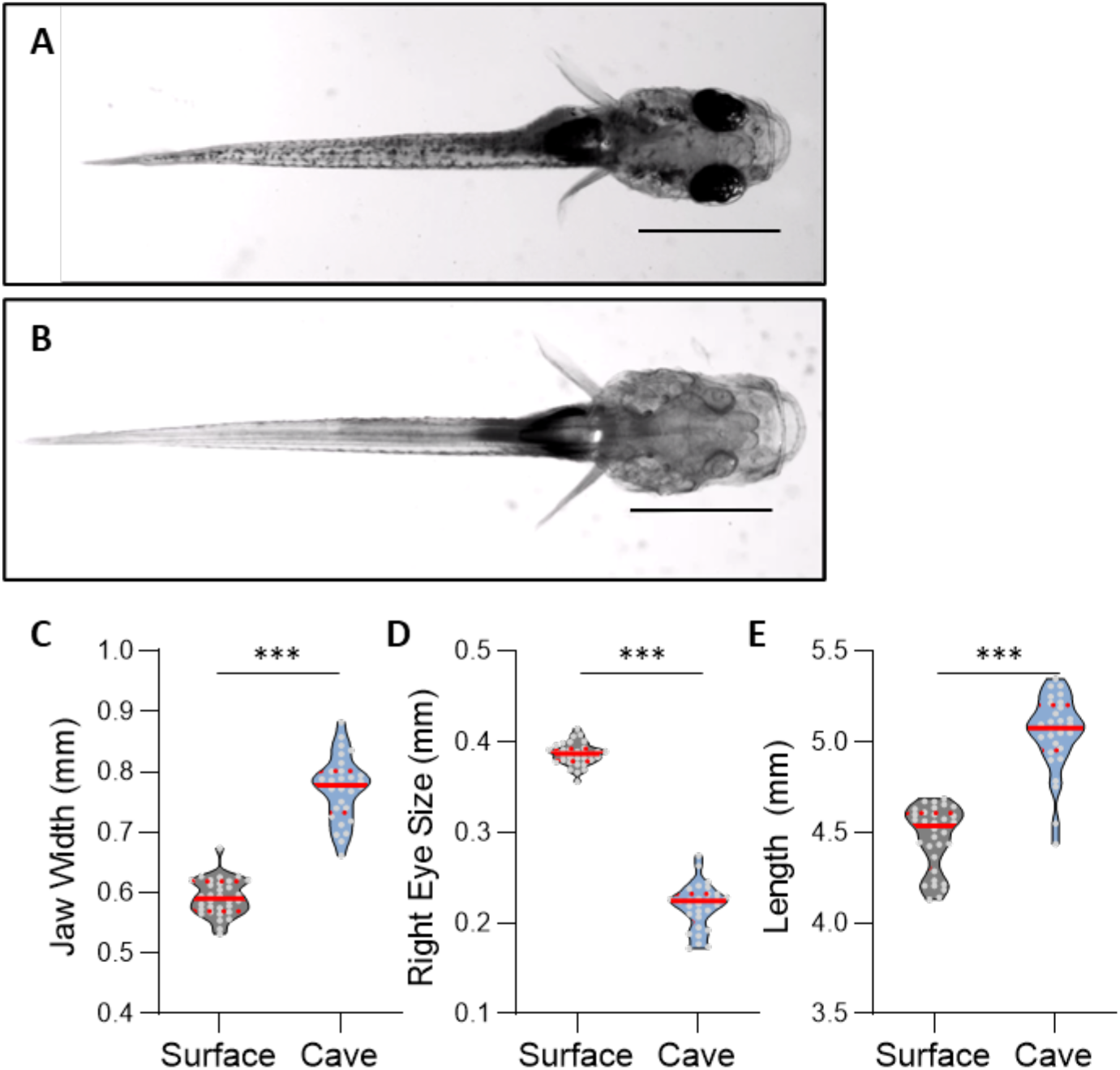
Morphological differences between purebred surface and Pachón cavefish populations. **A**,**B)** Dorsal image of surface **(A)** and Pachón cavefish **(B)**. Scale bars denote 1mm. **C)** Jaw width is significantly greater in cavefish compared to surface fish (t-test: t_59_=16.48, *P*<0.0001). **D)** Eye size is significantly greater in surface fish (t-test: t_59_=33.80, *P*<0.0001). **E)** Length is significantly greater in cavefish compared to surface fish (t-test: t_59_=11.55, *P*<0.0001). For each trait, the median (center line) as well as 25^th^ and 75^th^ percentiles (top and bottom lines) are shown. Circles represent values from individual fish. *** denotes P<0.001.

To define behavioral differences between surface and Pachon cavefish, we developed a behavioral analysis pipeline to quantify ecologically relevant behaviors in succession, assessing startle response kinematics, followed by prey capture, and finally locomotor activity in the same individual fish. We focused on behaviors related to foraging and predator evasion, as food scarcity and reduced predation are major changes associated with the cave environment (Elliott 2016; McGaugh *et al*. 2020). To measure feeding behavior, we recorded the response of surface and cavefish during *Artemia* feeding and quantified the angle and distance of prey capture, two kinematic components that differ between visually and non-visually-related feeding (Figure 2A; Lloyd et al. 2018; Jaggard et al. 2020). Consistent with previous reports, strike angle was significantly greater in cavefish (Figure 2B; Lloyd et al. 2018). We also found the strike distance was reduced in Pachón cavefish compared to surface fish (Figure 2C). To assess escape response kinematics, plates containing fish in individual wells were fastened to a small vibration excitor and the response to escape-inducing vibration was measured with a high-speed camera (Figure 2D). The angular speed of cavefish was reduced, approaching significance (*P*=0.06), while the peak angle was also reduced (Figure 2E,F), consistent with a previous report that the escape response is blunted in cavefish (Paz *et al*. 2020). Finally, we measured locomotor activity in cavefish because it is a critical for aspects of predator avoidance and foraging. Cavefish are more active, presumably to allow increased foraging activity and exhibit increased wall-following behavior (Duboué *et al*. 2011). We quantified total locomotor activity and time spent in the center of the arena over a one-hour assay (Figure 2G). Pachón cavefish spent a shorter duration of time in in the center of the test arena compared to surface fish and exhibited a greater total amount of activity (Figure 2H,I). Together, these findings are consistent with previously published reports revealing robust differences in sensory and foraging behavior. The establishment of these phenotypes in 6 dpf fish tested in succession for each behavior provides an assay for examining inter-individual variability.

**Figure 2.** Behavioral variation in surface and cavefish. **A)** Diagram of prey capture apparatus. Videos were used to extract strike angle (red) and strike distance (blue) between *Artemia* and the head of the fish. **B)** Strike angle is significantly greater in cavefish than surface fish (t-test: t_41_=3.006, *P*<0.0045). **C)** Strike distance in surface fish is significantly greater than in cavefish (t-test: t_41_=2.209, *P*<0.0328). **D)** Image of startle reflex set up. Plate sits on a mini-shaker (black) to induce a startle. Videos were used to extract angular speed and peak angle (red). Grey arrow denotes head orientation at the initiation of the startle stimulus. **E)** Angular speed in surface fish is significantly greater than in cavefish (t-test: t_27_=3.629, *P*<0.0012). **F)** Peak angle in surface fish is greater, approaching significance, than cavefish (t-test: t_27_=1.928, *P*<0.0645). **G)** Image of locomotor assay where fish were recorded in individual wells for 1-hr and to analyze total locomotor activity and time in the center of the well (grey area). **H)** Time spent in the center in surface fish is significantly greater than cavefish (t-test: t_37_=4.710, *P*<0.0013) **I)** Total distance in surface fish is significantly less than in cavefish. (t-test: t_37_=6.506, *P*<0.0001). For each trait, the median (center line) as well as 25^th^ and 75^th^ percentiles (dotted lines) are shown. Circles represent values from individual fish. * denotes *P*<0.05, ** denotes *P*<0.01; *** denotes *P*<0.001.

To examine the relationship between traits in surface and Pachón cavefish, we first examined the correlation between morphological traits. For most morphological traits there was a strong correlation between eye size, head width, length, and jaw width (Figure S4A,B). Conversely, there were far fewer significant interactions between individual components of behavior (Figure S4C,D). Surprisingly for cavefish, total locomotor activity significantly associated with peak angle, and angular speed, revealing a relationship between locomotor behavior and startle reflex (Figure S4C). In addition, in both surface and Pachón cavefish, angular speed associated with peak angle, suggesting a relationship between both metrics of startle reflex (Figure S4C,D). Finally, we examined the relationship between morphological and behavioral traits (Figure S4E,F). There were far fewer correlations between behavioral and morphological traits, than for morphology alone. In cavefish, total distance associated with many aspects of size, yet this was not observed in surface fish (Figure S4E,F). Taken together, these findings reveal strong associations between morphological traits and fewer between individual behaviors in pure populations of surface and cavefish.

We sought to define whether any of the morphological differences identified between surface fish and cavefish genetically segregate, a phenotype that would suggest they are governed by shared genetic architecture. To examine the relationship between different morphological traits, including body sizes, eyes, and pigmentation, we generated F2 surface-cave hybrids by crossing F1 offspring of the pure-bred surface fish and Pachón cavefish (Figure 3A). We were not able to detect intermediate pigmentation levels and this trait was scored in a binary fashion that includes fish that are heterozygous and homozygous for surface fish *oca2*. We quantified individual fish for numerous traits including eye size, length, height, and jaw size (Figure 3C,D,E). First, we quantified whether there was an interaction between these traits and pigmentation. Across all morphological traits measured in F2 offspring, there were no significant differences between pigmented and albino fish (Table 1 and Figure S5A-G). We directly compared eye size in albino and pigmented fish and found no relationship, suggesting the albinism gene *oca2* is not involved in the development of eye size (Figure 3C). We performed a Spearman’s Rank Correlation Coefficient across all variables and found traits related to size had strong associations with one another (Figure 3B). These studies revealed that the majority of morphological traits were linked, including an association between jaw width and overall size. While the difference did not reach significance, albino fish trended towards having larger eyes (*P*=0.06; Figure 3C). However, when eye-size was corrected for body length, eye size was larger in pigmented hybrids than albino hybrids (*P*<0.05; Figure S5H), raising the possibility that shared genes, or closely linked genes, contribute to both processes. It has previously been suggested that jaw width is associated with eye-size (Yamamoto *et al*. 2009) and we observed a significant interaction between these traits (Figure 3D). We also observed a significant interaction between jaw width and overall head width, suggesting jaw width is likely specified by the overall head size of the animal (Figure 3E). Together, these findings suggest the overall growth rate of cavefish is accelerated and the genes regulating different features of growth co-segregate in cavefish.

**Figure 3.**
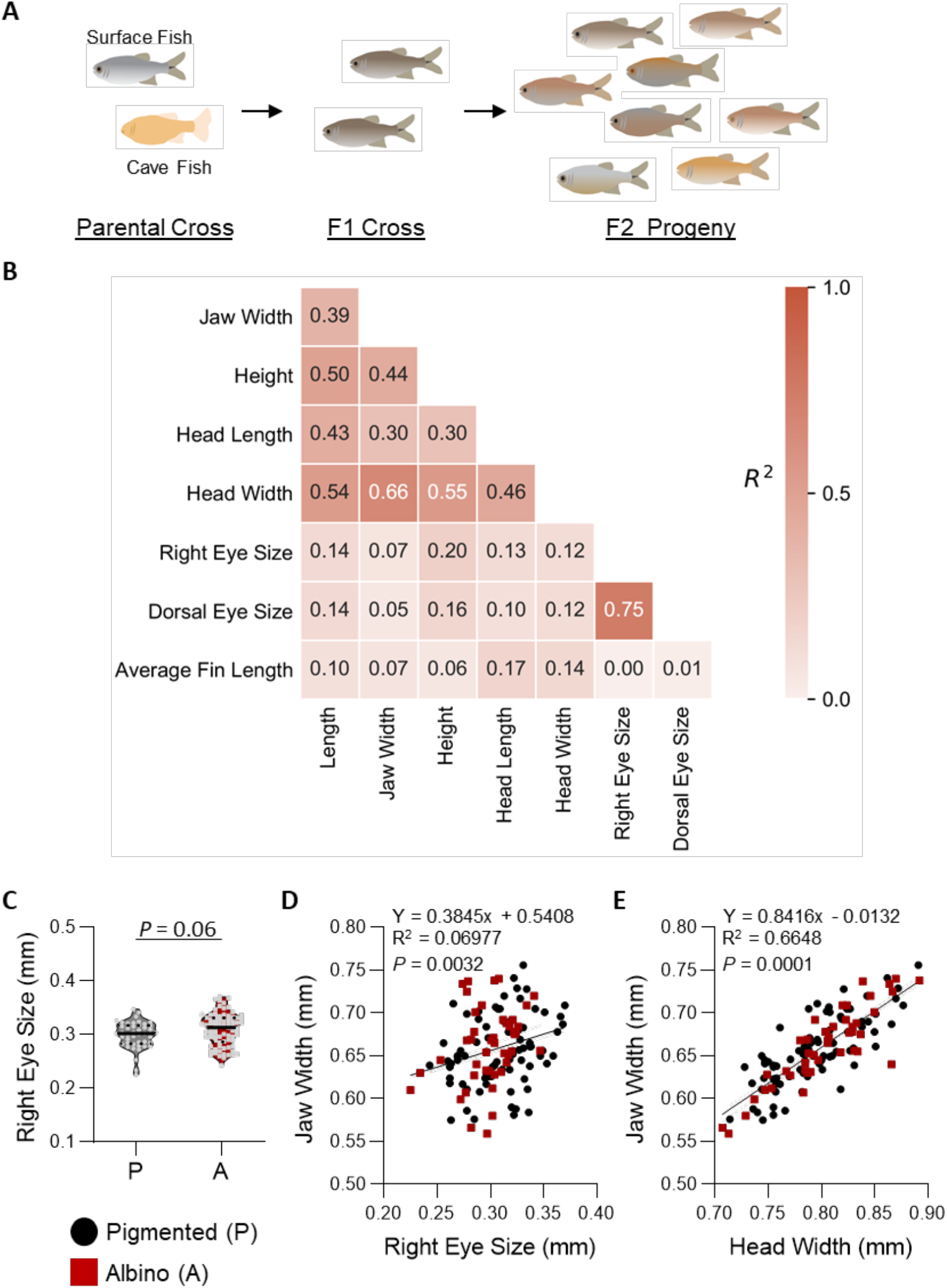
Morphological traits in F2 hybrid offspring. **A)** Cross-breeding process between purebred Surface (silver) and Pachón (albino) to produce F1 progeny, and F1 crosses to produce F2 hybrids used for study. **B)** Heat map of the correlations between morphological traits in F2 offspring (R^2^ values shown). **C)** Eye size does not differ between pigmented and albino individuals (t-test: t_121_=1.856, *P*<0.0659). **D)** Linear regression between jaw width and eye size reveals a significant association (F_1,121_=9.076, *P*<0.0032). **E)** Linear regression between head width and jaw width reveals a significant association (F_1,121_=137.5, *P*<0.0001). Albino individuals are depicted as red squares, while pigmented individuals are depicted as black circles.

It is possible that the many behavioral differences in cavefish evolved independently of one another, or that they are governed by shared genetic architecture. To test the relationships between these traits, we measured the behavior of individual F2 hybrids for locomotor behavior, prey-capture, and escape reflex (Figure 4A). We then performed rank-correlation analysis across all behavioral traits (Figure 4B). We identified a correlation between angular speed and peak angle in escape responses, but not with other behavioral variables tested (Figure 4C). Additionally, no significant associations were identified for variables of prey capture and escape reflexes, suggesting the evolved differences for each behavior in cavefish occurred through independent genetic mechanisms (Figure 4D). Together, these findings suggest there is little shared genetic or functional relationship between three behaviors that are thought to be critical to cave adaptation.

**Figure 4.**
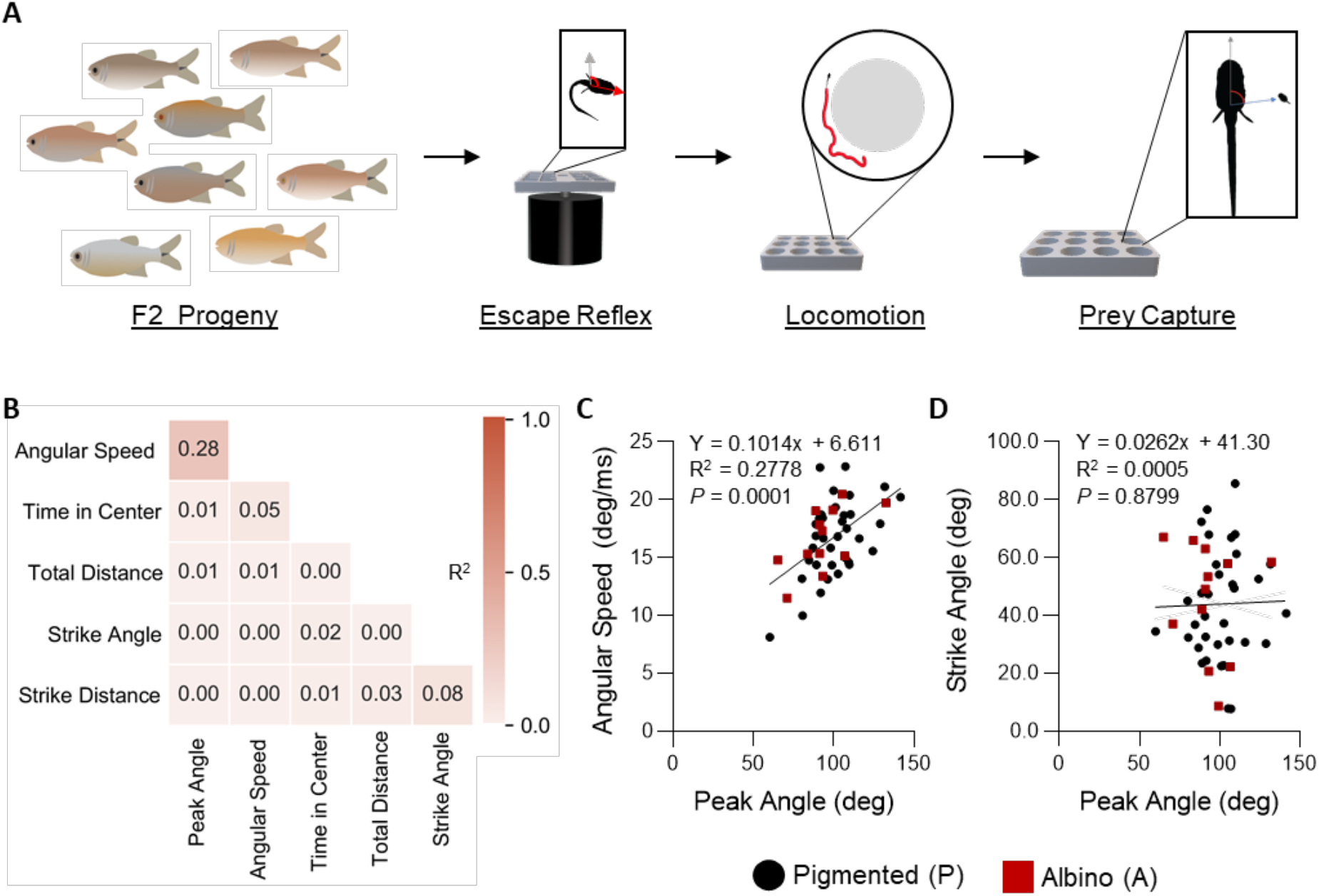
Relationship between behavioral traits within F2 hybrids. **A)** Schematic for behavioral analysis where individual F2 fish were tested for locomotor activity, escape reflex, and then prey capture in succession. **B)** Heat map of the correlations between behavioral traits in F2 offspring (R^2^values shown). **C)** Peak angle of the escape reflex is associated with angular speed (F_1,45_=17.31, *P*<0.0001). **D)** Peak angle during the escape reflex is not associated with feeding strike angle (F_1,44_=0.0231, *P*<0.8799). Albino individuals are depicted as red squares, while pigmented individuals are depicted as black circles.

Numerous studies have revealed associations between morphological and behavioral evolution (Alié *et al*. 2018). To examine the possibility that the behaviors studied in our analysis pipeline relate to anatomical changes, we compared the associations between anatomical and behavioral traits measured in F2 offspring (Figure 5A). We identified significant negative correlations between angular speed in escape response and body length as well as between peak angle in escape response and head length, suggesting a trade-off between increased size and reduced escape response performance in cavefish (Figure 5D). We also identified an association between albinism and total swimming distance, consistent with the notion that mutations in *oca2* confer sleep loss and altered locomotor activity (Figure 5B; Bilandžija et al. 2018; O’Gorman et al. 2020). These differences are likely not reflective of general locomotor abnormalities in albino hybrids because the time in the center did not differ across each population (Fig 5C). Taken together, these findings suggest the evolution of behavioral and morphological phenotypes are largely governed by independent genetic architecture.

**Figure 5.**
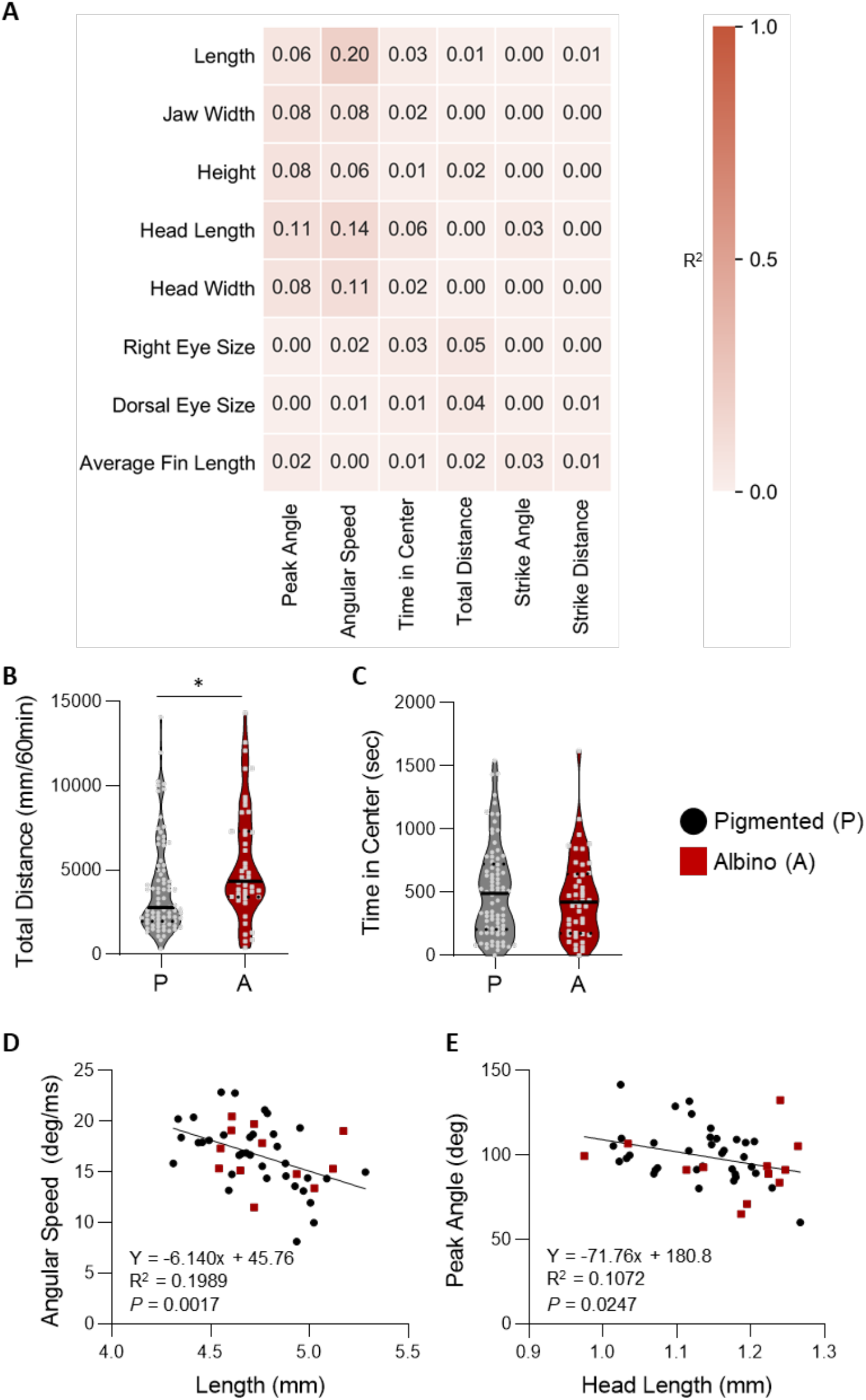
Comparison between morphological and behavioral traits within F2 hybrid offspring. **A)** Heat map of the correlations between morphological and behavioral traits in F2 offspring (R^2^ values shown). **B)** Total distance is significantly greater in albino (A) than in pigmented (P) individuals (t-test: t_125_=2.636, *P*<0.0095). **C)** Time in the center does not differ between pigmented and albino individuals (t-test: t_125_=0.8155, *P*<0.4163). For each trait, the median (center line) as well as 25^th^ and 75^th^ percentiles (dotted lines) are shown. **D)** Standard length and angular speed are significantly correlated in F2 hybrid individuals (R^2^=0.1989). **E)** There is a significant association between peak angle and head length in F2 individuals (R^2^=0.1072). Albino individuals are depicted as red squares, while pigmented individuals are depicted as black circles. ** denotes *P*<0.01.

## Discussion

Across taxa, environmental perturbation leads to the evolution of many behavioral and morphological traits (Stern 2013). Patterns of genetic covariations can influence the rate and direction of phenotypic evolution as seen in the three-spine stickleback populations, resulting in changes in body armor, aggression, and social behaviors (Leinonen *et al*. 2011; Peichel and Marques 2017). Similarly, rapid evolution of species in East African cichlids has led to dramatic changes in many traits including coloration, craniofacial morphology, aggression, and locomotor behavior (Kocher 2004; Powder and Albertson 2016; Salzburger 2018□). Understanding how defined ecological factors impact the evolution of these traits, as well as their genetic bases, is a central question in evolution. Examining numerous behavioral and morphological traits in hybrids with robust evolutionarily-derived differences can be applied to many different species to examine the genetic basis of trait evolution, and whether traits are governed by shared genetic architecture.

To investigate the relationship between genetic architecture underlying the evolution of behavioral and morphological traits, we quantified segregation of these traits in surface-Pachón cave F2 hybrids. The approach of examining numerous behavioral and morphological traits in hybrids with robust evolutionarily-derived differences can be applied to a broad number of species to examine the genetic basis of trait evolution, and whether traits are governed by shared genetic architecture.

We performed broad analyses of morphological and behavioral traits that suggest the genetic architecture underlying multiple aspects of increased body size including head size, head width and body length, and jaw width are all related. This suggests defined genetic changes have led to an overall increase in growth rate during early development in cavefish. However, there were no associations between eye size and albinism with any other morphological traits. These findings are surprising because QTL analysis has found that eye size and albinism localize to overlapping QTL, raising the possibility that albinism is pleiotropic (Protas *et al*. 2008). Further, increased jaw size has been previously associated with a reduction in eye size(Yamamoto *et al*. 2009). Therefore, increasing the sample size or testing at different developmental stages may identify additional associations between traits including developmentally-specified associations, with interactions at some stages, but not early development. Alternatively, the effect size of many traits may be small and were missed during this study. It is important to note that we only examined superficial morphological traits, and it is possible that a more detailed analysis would uncover additional traits that are related. For example, cavefish have differences in craniofacial morphology, tooth development, and an expansion of lateral line neuromasts (Varatharasan *et al*. 2009; Yoshizawa *et al*. 2010; Gross *et al*. 2014; Atukorala and Franz-Odendaal 2018). In addition, we did not test possible interactions between morphological traits and brain neuroanatomy that have previously been reported (Jaggard *et al*. 2019; Loomis *et al*. 2019). Therefore, it is possible that a more detailed analysis will reveal broader genetic interactions.

Most behavioral traits studied here are unlinked from the observed anatomical changes. We did, however, identify an association between increased length and reduced angular speed during the escape response, revealing a potential trade-off between size and escape performance in fish. While limited information is available about the relationship between body size and escape response in larvae, in adult damsel fish size is correlated with increased escape velocity (McCormick *et al*. 2019). Given that cavefish appear to lack macroscopic predators (Jeffery 2008; Kowalko 2020), it is possible that rapid growth is advantageous in order to develop resistance to starvation, even at the expense of reduced escape abilities. Broadly, we found that pigmentation did not associate with nearly all traits tested (with the exception of eye size), suggesting that loss of *oca2* function is relatively specific to albinism and does not impact the behaviors tested. This is surprising given the role of *oca2* as in monoamine function. In *oca2* mutants, norepinephrine and dopamine levels are elevated, and these neurotransmitters are linked to many behaviors including foraging (Bilandzija *et al*. 2013; Bilandžija *et al*. 2018). We did observe increased locomotor behavior in albino mutants, consistent with previous findings that sleep is reduced F2 surface x cave hybrids or surface fish with engineered mutations in *oca2* (O’Gorman *et al*. 2020). Therefore, our findings suggest that the differences in behavioral and morphological traits examined here largely evolved through independent genetic mechanisms, though there are likely to be trade-offs between body size and escape behavior during early development.

A central question in the field relates to the ecological factors that drive many of the evolved differences in behavior and morphology in cavefish. Here, we confirm previous findings revealing sensory-motor changes in prey capture and startle response, as well as changes in locomotion (Lloyd *et al*. 2018; Jaggard *et al*. 2020). It is unclear which aspects of cave ecology are likely to drive these differences. Caves and surface habitats differ in many ways, including constant darkness, which is proposed to underlie increased dependence on the lateral line during feeding behavior (Yoshizawa *et al*. 2010, 2012). There is also speculation that the caves are nutrient poor compared to surface environments, and this underlies the evolution of sleep loss, however this has not been investigated systematically in the cave environment (Krishnan and Rohner 2017). It is possible that the increased jaw size in cavefish is related to the size of the prey consumed by juveniles, allowing for larger prey, greater suction during feeding, or improved success during lateral-line dependent feeding that involves lateral movement during capture (Yoshizawa *et al*. 2010; Holzman *et al*. 2014; Lloyd *et al*. 2018). Conversely, it has previously been shown that enhanced prey capture abilities of larval cavefish are independent from eye loss (Espinasa *et al*. 2014). Little is known of the foraging behavior of fish in natural conditions, especially at the larval stage. The stomach contents of adult fish, identifying a diet of arthropods and there is speculation that cavefish consume bat guanos deposited from bat colonies that inhabit the majority of the caves (Espinasa *et al*. 2017). Further investigation of the abiotic and biotic ecology of the caves are likely to contribute to our understanding of evolution, and comparisons of surface and cavefish across different developmental stages should improve our understanding of how the studied traits have evolved.

We examined hybrids of thesurface and Pachón cave populations. We chose this population because geological, genomic, morphological evidence suggests the Pachón population is one of the most troglomorphic, and therefore the most commonly used in Mexican tetra studies. The largely independent evolution of at least 30 different cave populations offers a unique opportunity to study the evolution of various traits. Shared genetic changes underlie evolution in a number of populations. For example, different mutations in the pigmentation gene *oca2* directly lead to albinism in the Molino and Pachón populations (Protas et al 2008). In addition, complementation analysis between independently evolved cavefish populations suggest different genetic changes underlie eye loss in the Pachón and Molino populations (Sifuentes-Romero *et al*. 2020). However, the presence of convergent evolution in these populations increases the likelihood of differences in traits among individual populations. For example, sleep loss in the Pachón population is dependent on enhanced lateral line function, while sleep loss in Tinaja and Molino fish is independent of the lateral line (Jaggard *et al*. 2017). Therefore, the systematic relationship between evolved traits across multiple independently evolved populations of cavefish has potential to uncover whether shared principles governed repeated evolution following similar ecological changes.

In this study we exclusively examined behavior at 6 days post fertilization. This is an age typically used in zebrafish for genetic manipulations including performing whole brain imaging, and a recently developed neuroanatomical atlas in *A. mexicanus* compared different populations at this age (Halpern *et al*. 2008; Keene and Appelbaum 2019; Jaggard *et al*. 2020). Further, hybrid analysis studies often require large numbers of fish, and testing fish at 7dpf is much more accessible. While the differences between surface fish and cavefish behavior for sleep, foraging, and wall-following (or reduced time in center) are similar in 7 dpf fish and adults, there may be developmentally-specified effects. For example, at 30 dpf the prey capture distance (distance between prey and fish at the start of the attack) is greater in surface fish than cavefish (Lloyd *et al*. 2018), however, we report that here it is reduced at this timepoint at 6 dpf. Despite these differences, multiple studies now confirm that the attack angle is greater in cavefish as early as 7 dpf through 30 dpf. Therefore, the phenotypes observed may vary across developmental, and therefore any genetic relationships identified through the approach used here may not generalize across development. In addition to the behaviors examined, the behaviors of adults are thought to be more complex, and therefore may allow for more detailed analysis of the relationship between differentially-evolved behaviors. Many behavioral differences have only been described in adults including schooling, aggression, vocalizations, and vibration attraction, and therefore could not be included in the analysis applied here. The approach of generating a pipeline for examining trait interactions could be applied to adult animals allowing for investigation of the interactions between a broader number of traits.

This investigation sought to understand the relationship between many different evolved traits in F2 surface x cave hybrid fish. While our analysis was limited to phenotyping, previous studies have performed mapping studies to localize genomic regions associated with numerous traits including albinism, locomotor behavior, eye size, social behavior, nonvisual sensory systems (O’Quin and McGaugh 2016). Sequenced genomes for Pachón cave and surface populations of *A. mexicanus* are available and can be applied for mapping traits observed (McGaugh *et al*. 2014; Warren *et al*. 2021). The behavioral pipeline approach used in this study would be particularly powerful because it would allow for genomic mapping of many traits in a relatively small number of animals. In addition to genomic approaches, gene-editing approaches have been applied to functionally validate data obtained from genomic mapping or transcriptional analysis, revealing potential for this approach to identify novel genetic regulators of many different cave evolved traits (Ma *et al*. 2015; Klaassen *et al*. 2018; Stahl *et al*. 2019). Finally, the approach used and its potential for mapping is not limited to *A. mexicanus*. Hybrid analysis and mapping is widely used in other fish species and applying a behavioral pipeline to identify genetic architecture associated with trait evolution has potential for identifying genes in many different models of evolution.

## Materials and Methods

### Fish Husbandry

Animal husbandry was carried out as previously described (Stahl *et al*. 2019) and all protocols were approved by the IACUC Florida Atlantic University. Fish were housed in the Florida Atlantic University core facilities at 23 °C± 1 °C constant water temperature throughout rearing for behavior experiments. Lights were kept on a 14:10 h light-dark cycle that remained constant throughout the animal’s lifetime. Light intensity was kept between 25 and 40 lx for both rearing and behavior experiments. Adult fish were fed a diet of black worms to satiation twice daily at zeitgeber time (ZT) 2 and ZT12, (Aquatic Foods, Fresno, CA,) and standard flake fish food during periods when fish were not being used for breeding (Tetramine Pro). All fry used for experiments were reared on live *Artemia* beginning at 4dfp and fed twice daily through the end of experiments at 7 dpf.

### Behavioral analysis pipeline

All fish tested, including surface fish, Pachón cavefish, and surface x cave F2 hybrids followed the same behavioral analysis pipeline. At 6 dpf fish were removed from bowls, transferred to plates and tested for startle reflex as described below. All tests took place between ZT0 and ZT4 in the light. Following completion of startle response assays, larvae were returned to their well plates and transferred to the locomotor assay. Locomotion behavior experiments were run as 4 trials a day, with a 60 min trial interval. Behavioral videos were captured between ZT4 and ZT8. Immediately following locomotor assays at ZT8, larvae were returned to their well plates and then transferred to prey capture arenas containing *Artemia* nauplii, where their behavior was recorded for a single period of 30 minutes. After this period, larvae were returned to their original well plates and incubator overnight. The following day, larvae were retrieved and photographed for measurement of morphological traits. The detailed of each part of the procedure are described below. At all stages care was taken to avoid mixing u individual fish throughout the process.

### Prey capture

Prey capture behavior was recorded as previously described, with minor modifications, described below (Lloyd et al, 2018). Video was acquired using a USB 3.0 camera (LifeCam Studio, Microsoft) fitted with a zoom lens (75 mm DG Series Fixed Focal Length Lens, Edmund Optics Worldwide), and recorded with VirtualDub2 (v44282). All images were acquired at 30 frames per second. Recording chambers were illuminated with custom-designed infrared LED source (Infrared (IR) 850 nm 5050 LED Strip Light, Environmental Lights). All recordings were performed in 6 dpf fry from zeitgeber (ZT) 8 to ZT10. For larval fish recordings, individual fish were placed in 24 well tissue culture plates (Cellvis) or custom-made chambers, filled with ∼3 mm of water to constrict the larvae to a single focal plane. Fish were allowed to acclimate for 2 min prior to the start of the experiment. To record feeding behavior, approximately 30 *Artemia nauplii* were added to each well and fish were imaged for 30 minutes.

Recordings were analyzed using ImageJ 1.52a (National Institutes of Health; Bethesda, MD). Chamber diameter was set using ImageJ’s native “Set Scale” function, and strike distance and angle were measured for all successful feeding events, using ImageJ’s “Line” and “Angle” tools. Measurements of both strike distance and angle were taken in the frame prior to initiation of movement towards the prey. Strike distance was defined as the shortest distance between the edge of the fish’s body and the prey. Strike angle was defined as the angle between a line extending down the fish’s midline, terminating parallel with the pectoral fins, and a line extending from this point to the center of the prey. Measurements of each strike were averaged to calculate the mean strike distance and angle for that individual, and any recording with fewer than three feeding events was excluded from analysis.

### Startle response

We assessed startle response probability and kinematics as previously described (Paz et al, 2019). Assays were conducted in a temperature-controlled environment maintained at 24°C. Individual F2, surface, or Pachon larvae were placed in square wells on a custom 3D printed polyactic acid 16-well plate, which was mounted onto a vertically-oriented vibration Exciter controlled by a multi-function I/O device and custom Labview 2018 v.18.0f2 (National Instruments, Austin, TX) scripts. To optimize the quality of video recordings, only 8 wells were used at a time. Each assay consisted of an initial 10 minute acclimation period followed by six 500 Hz square wave stimuli of 50 ms duration with a 10 minute interstimulus interval, resulting in a total duration of one hour per assay. A total of 128 F2 larvae were assayed. An LED was connected directly to the signal driving the exciter so that its flashing could be used to identify the start and end of each stimulus in video recordings. C-start responses were identified as accelerated, simultaneous flexion of the head and tail in the same direction. Response probability is reported as the total number of c-starts performed by a larva divided by the total number of stimuli to which the larva was exposed (six). Beginning from the frame immediately preceding the stimulus start (as indicated by the LED turning on), the “angle” tool on ImageJ 1.52a (National Institutes of Health; Bethesda, MD) was used to determine the change in orientation of the larvae over the course of the stimulus, and these measurements were used to determine response latency, angular speed, and peak angle. Response latency is defined as the time interval between stimulus onset and a change in orientation of at least 10 degrees.

### Locomotor behavior

We measured locomotion activity as previously described (Jaggard JB, 2019). Videos were captured using a Basler ace ac1300-200 um USB 3.0 digital camera (Edmund Optics Inc., NJ; CAT#33978) with a 16mm C series lens (Edmund Optics Inc., CAT#67714) and a UV-VIS filter (Edmund Optics Inc., CAT#65716). *Astyanax mexicanus* populations were placed in single wells of a 6-well plate (34.8 mm diameter; #S3506, Corning Inc., Corning, NY), acclimated for 10 minutes and recorded for one hour. Distance, velocity, and time spent in the center and border were tracked and analyzed using the software EthoVision (Ethovision X14, Noldus Information Technology, Wageningen, NLD). Raw data was binned and transformed using custom made MATLAB scripts (available on request).

### Morphological analysis

Pure surface and Pachón cavefish or F2 offspring were anesthetized in 0.1 M Tricaine at 7 days post-fertilization for imaging. Each fish was imaged both dorsally and laterally. Images were standardized against a 1mm measurement and were uploaded to Fiji ImageJ (Schindelin *et al*. 2012). The 1mm measurement was measured by the line variable in ImageJ and established as a global calibration for all future measurements. Each fish image was used to measure lengths for the following: standard length, head depth, eye size lateral, eye size dorsal, jaw width, head width, head length, fin length left and right (which were averaged to a single value). Standard length measurement connected the tip of the upper lip to the end of the tail at its greatest length. Head depth was measured as the length from the dorsal edge of the head to the ventral edge of the head. Eye size lateral was determined as the length between either side of the widest part of the right eye (under microscope when fish is placed laterally). The remaining measurements were done from a dorsal perspective. Eye size dorsal was measured from the outside of the widest part of the eye from the center edge to the lateral edge. Jaw width was measured as the length of the widest part of the head rostral to the eyes. Head width was measured as the widest length of the head caudal to the eyes. Head length was recorded as the length from the tip of the upper lip to the end of the head, established when it meets the swimmer bladder. Finally, fin lengths were measured as the greatest length where each fin attaches to the head to the tip of the end of each fin. Every measurement was completed by two different raters to determine inter-rater reliability, and both measurements were averaged to find final values for analysis.

### Statistical analysis

All morphological and behavioral traits are presented as violin plots; indicating the median, 25^th^, and 75^th^ percentiles. All statistical analyses were performed using Instat software (Graphpad Prism 8.4.3). For each trait, normality was assessed visually from a QQ plot and then a parametric t-test was performed. To visualize the relationship between two traits, a linear regression was performed. R^2^ heatmaps were generated using python’s SciPy and Seaborn modules. R^2^ values were obtained by performing an independent linear regression on each pair of variables.

## Acknowledgements

The authors are grateful to Julia Pasquale for assistance with behavioral analysis, as well as Peter Lewis and Arthur Lopatto for assistance with fish care. This work was supported by an NIGMS U-RISE award (T34GM136486) to ACK an NIGMS R01 (1R01GM127872) to ACK and an NSF-EDGE award to JEK and ERD (1923372).

## Supplemental Figure Legends

**Supplementary Figure 1: Landmarks used for morphometric analysis**. Each line depicts a measurement used for morphometric analysis in surface fish (top) and Pachón cavefish (bottom). Both dorsal (right) and side views were imaged for each individual fish and used for quantification.

**Supplementary Figure 2: Quantification of anatomical differences between surface fish and Pachón cavefish. A)** Height is significantly greater in cavefish compared to surface fish (t-test: t_59_=6.710, *P*<0.0001). **B)** Head length is significantly greater in cavefish compared to surface fish (t-test: t_59_=14.65, *P*<0.0001). **C)** Head width is significantly greater in cavefish compared to surface fish (t-test: t_59_=15.37, *P*<0.0001) **D**) Dorsal eye size is significantly reduced in cavefish compared to surface (t-test: t_59_=36.49, *P*<0.0001). **E**) There is no difference in average fin length between surface fish and cavefish (t-test: t_59_=1.422, *P*<0.1603). For each trait, the median (center line) as well as 25^th^ and 75^th^ percentiles (dotted lines) are shown. Circles represent values from individual fish. *** denotes P<0.001.

**Supplementary Figure 3: Quantification of normalized anatomical differences between surface fish and Pachón cavefish. A)** Normalized jaw width is significantly greater in cavefish compared to surface fish (t-test: t_59_=11.26, *P*<0.0001). **B)** Normalized eye size is significantly reduced in cavefish compared to surface fish (t-test: t_59_=49.76, *P*<0.0001). **C)** Normalized height is significantly reduced in cavefish compared to surface (t-test: t_59_=6.971, *P*<0.0001). **D)** Normalized head length is significantly greater in cavefish compared to surface (t-test: t_59_=9.817, *P*<0.0001). **E)** Normalized head width is significantly greater in cavefish compared to surface fish (t-test: t_59_=7.634, *P*<0.0001). **F)** Normalized dorsal eye size is significantly reduced in cavefish compared to surface (t-test: t_59_=62.14, *P*<0.0001). **G)** Normalized average fin length is significant reduced in cavefish compared to surface (t-test: t_59_=6.563, *P*<0.0001). For each trait, the median (center line) as well as 25^th^ and 75^th^ percentiles (dotted lines) are shown. Circles represent values from individual fish. *** denotes P<0.001.

**Supplementary Figure 4: Pairwise correlations between traits in surface and cave fish**. For each pairwise comparison, R^2^ values are shown. **A-B)** Heat map of the correlations between morphological traits in **(A)** cave and **(B)** surface fish. **C-D)** Heat map of the correlations between behavioral traits in **(C)** cave and **(D)** surface fish. **E-F)** Heat map of the correlations between morphological and behavioral traits in **(E)** cave and **(F)** surface fish.

**Supplementary Figure 5: Quantification of anatomical differences between pigmented and albino F2 hybrid offspring. A)** Length does not differ between pigmented and albino individuals (t-test: t_121_=0.3415, *P*<0.7334). **B)** Jaw width does not differ between pigmented and albino individuals (t-test: t_121_=0.7447, *P*<0.4579). **C**) Height does not differ between pigmented and albino individuals (t-test: t_121_=0.8388, *P*<0.4032). **D)** Head length does not differ between pigmented and albino individuals (t-test: t_121_=0.3090, *P*<0.7578). **E**) Head width does not differ between pigmented and albino individuals (t-test: t_121_=1.265, *P*<0.2084). **F**) Dorsal eye size does not differ between pigmented and albino individuals (t-test: t_121_=1.780, *P*<0.0775). **G**) Average fin length does not differ between pigmented and albino individuals (t-test: t_121_=0.8878, *P*<0.3764). **H**) Normalized eye size is significantly reduced in albino compared to pigmented hybrid offspring (t-test: t_121_ =2.107, *P*<0.0372). For each trait, the median (center line) as well as 25^th^ and 75^th^ percentiles (dotted lines) are shown. Albino individuals are depicted as red squares, while pigmented individuals are depicted as black circles.

**Supplementary Figure 6: Quantification of behavioral differences between pigmented and albino F2 hybrid offspring. A**) Strike angle does not differ between pigmented and albino individuals (t-test: t_116_=0.0488, *P*<0.9611). **B)** Strike distance does not differ between pigmented and albino individuals (t-test: t_116_=0.0182, *P*<0.9855). **C)** Angular speed does not differ between pigmented and albino individuals (t-test: t_45_=0.0852, *P*<0.9325). **D)** Peak angle does not differ between pigmented and albino individuals (t-test: t_45_=1.354, *P*<0.1826). For each trait, the median (center line) as well as 25^th^ and 75^th^ percentiles (dotted lines) are shown. Albino individuals are depicted as red squares, while pigmented individuals are depicted as black circles.

